# Population density affects the outcome of competition in co-cultures of *Gardnerella* species isolated from the human vaginal microbiome

**DOI:** 10.1101/2021.02.09.430506

**Authors:** Salahuddin Khan, Janet E. Hill

**Author notes:** To whom correspondence should be addressed: 52 Campus Drive, Saskatoon, SK S7N 5B4, Canada, E-mail: SK, JEH.

## Abstract

Negative frequency-dependent selection is one possible mechanism for maintenance of rare species in communities, but the selective advantage of rare species may be checked at lower overall population densities where resources are abundant. *Gardnerella* spp. belonging to cpn60 subgroup D, are detected at low levels in vaginal microbiomes and are nutritional generalists relative to other more abundant *Gardnerella* spp., making them good candidates for negative frequency-dependent selection. The vaginal microbiome is a dynamic environment and the resulting changes in density of the microbiota may explain why subgroup D never gains dominance. To test this, we co-cultured subgroup D isolates with isolates from the more common and abundant subgroup C. Deep amplicon sequencing of rpoB was used to determine proportional abundance of each isolate at 0 h and 72 h in 152 co-cultures, and to calculate change in proportion. D isolates had a positive change in proportional abundance in most co-cultures regardless of initial proportion. Initial density affected the change in proportion of subgroup D isolates either positively or negatively depending on the particular isolates combined, suggesting that growth rate, population density and other intrinsic features of the isolates influenced the outcome. Our results demonstrate that population density is an important factor influencing the outcome of competition between *Gardnerella* spp. isolated from the human vaginal microbiome.

## Introduction

Negative frequency-dependent selection is an evolutionary phenomenon that maintains rare genotypes (variants) in complex communities and occurs when the relative fitness of a variant is higher when its relative abundance in the population is low [1, 2]. Resource competition is a major force that can selectively benefit rare variants, contributing to negative frequency-dependent selection. Slower growing variants that can utilize a wide range of nutrients (nutritional generalists) have a selective advantage over faster growing nutritional specialists once the availability of nutrients favoured by the specialists becomes limiting [3]. In a closed system, negative frequency-dependent selection could lead to the initially rare variant increasing in relative abundance in the population. In natural microbial communities, however, this process may be checked by factors including the density of the population and the carrying capacity of the environment since the benefits of negative frequency-dependent selection only come into play when the more abundant members of the community experience a reduction in fitness [4]. Variants subject to negative frequency-dependent selection may be prevented from gaining dominance due to environmental changes that introduce a fresh supply of nutrients for the specialists or reduce overall population density, shifting the selective advantage to the more common variants.

We have shown previously that resource-dependent, scramble competition is prevalent among *Gardnerella* spp. isolated from the human vaginal microbiome [5]. Women with *Gardnerella* dominated vaginal microbiomes are usually colonized by multiple species, with *G. vaginalis* (cpn60 subgroup C), *G. leopoldii* and *G. swidsinskii* (subgroup A) being the most common and abundant [6, 7]. *G. piotii* (subgroup B) and strains belonging to several currently unnamed genome species (subgroup D) are less frequently detected and have not been observed to dominate the microbiome [6, 7]. In *in vitro* co-culture experiments, isolates from subgroups A, B and C had reduced fitness when co-cultured with isolates from other subgroups, but the growth rates of subgroup D isolates increased with increasing number of competitors [5]. Subgroup D isolates were subsequently shown to have a nutritional generalist lifestyle and limited niche overlap with other *Gardnerella* spp. suggesting that these isolates persist in the vaginal microbiome through negative frequency-dependent selection [8]. Vaginal microbiomes dominated by subgroup D, however, have not been reported [6, 7]. While this potentially an argument against negative frequency-dependent selection, another possibility is that the negative frequency-dependent selection of rare *Gardnerella* spp. is density-dependent. Vaginal dynamics affecting bacterial population density: turnover of epithelial cells, bacterial mortality rates, flow of vaginal fluid, changing niche capacity, and interactions of microbial species in the vaginal ecosystem, can reshuffle the densities of competing species [4, 9, 10]. As the vaginal ecosystem changes, the advantage that slow growing, generalist species have over others may be diminished because at lower population densities, the supply of nutrients available to fast-growing bacteria will plentiful and thus they will out-compete slower growing species even if they are generalists (Fig 1) [4].

**Fig 1.**
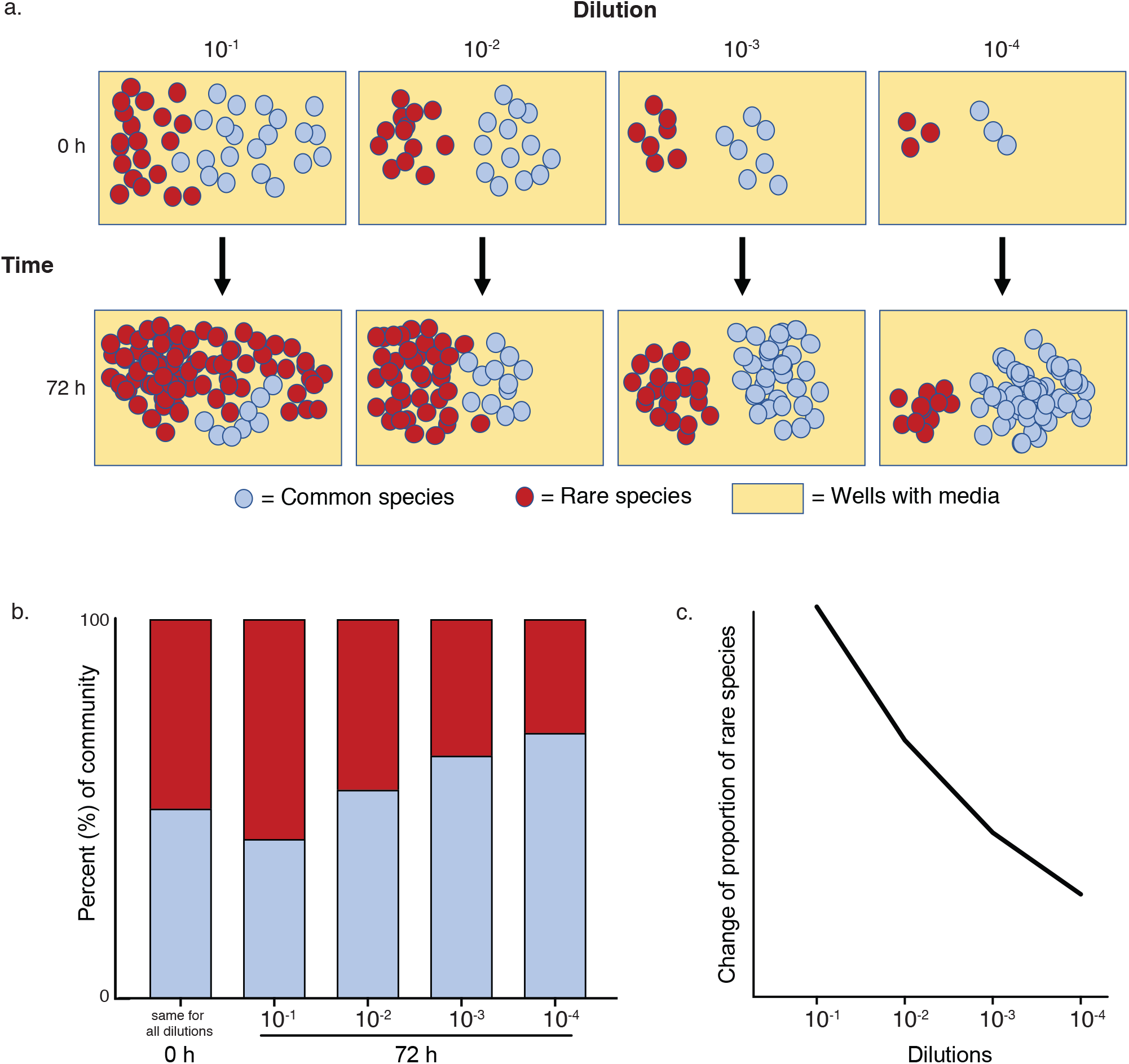
Effect of density on negative frequency-dependent selection. (a) If a common and a rare species are mixed at equal proportion at varying densities and grown for a fixed period of time, the selective advantage of the rare species will decrease with lower initial population densities. (b) The proportional abundance outcomes of the experiment shown in (a). (c) Using the initial and final proportional abundances of each isolate at varying densities, change of proportion is calculated. The change of proportion value of a species subject to density-dependent negative frequency-dependent selection will decrease with decreasing initial population density.

Demonstrations of the influence of density on negative frequency-dependent selection *in vitro* have been achieved by monitoring population dynamics of contrived microbial communities over a range of dilutions to mimic the dynamics of a real-world environment [3, 4]. The objective of our current study was to apply this approach to determine if population density affects the outcome of competitions involving *Gardnerella* subgroup D.

## Methods

### Bacterial isolates

Nine *Gardnerella* isolates (Subgroup C, *n* = 2, NR001, NR038; subgroup D, *n* = 7, NR002, NR003, NR043, NR044, NR047, WP012, N160) were used for this experiment. All isolates were revived from −80°C stocks by streaking on Columbia agar plates containing 5% (v/v) defibrinated sheep blood. For broth cultures, up to 10 isolated colonies from blood agar plates were picked and sub-cultured in brain heart infusion (BHI) broth supplemented with 10% (v/v) heat inactivated horse serum and 0.25% maltose (w/v) and incubated anaerobically for 18h (BD GasPak EZ Anaerobe Gas Generating Pouch System, NJ, USA).

### Co-culture experiment

Freshly grown broth cultures were adjusted to an OD of 0.5, and a loopful of each was streaked on Columbia blood agar to confirm viability. For each co-culture, an equal volume (250 μl) of one subgroup C and one subgroup D culture were mixed in 4.5 ml of BHI supplemented with 0.25% maltose to make a 10^−1^ dilution. Ten-fold dilutions up to 10^−4^ were made. Each of the two C isolates: NR001(C1) and NR038 (C2) were mixed with each of the seven D isolates: NR002 (D1), NR043 (D2), NR044 (D3), NR003(D4), N160 (D5), NR047 (D6), and WP012 (D7) resulting in 14 combinations (set 1: C1D1, C1D2, C1D3, C1D4, C1D5, C1D6, C1D7; and set 2: C2D1, C2D2, C2D3, C2D4, C2D5, C2D6, C2D7). Triplicate aliquots of 200 μl from each dilution of every combination were transferred into individual wells of 96-wells plates, and an additional aliquot was immediately pelleted and stored at −80°C (0 h). The total number of co-cultures of subgroup C and D thus created in 96-well plates was 168 (14 combinations × 4 dilutions × 3 technical replicates = 168). All co-cultures were incubated anaerobically for 72 h and OD_595_ was measured at 48 h and 72 h.

All C and D isolates were also grown alone in eight wells each as technical replicates. The OD_595_ of the monocultures were measured at 24 h, 48 h and 72h. To quantify planktonic growth of the monocultures, 72 h culture supernatant was transferred to a fresh 96-well plate and the OD_595_ was measured. The adhered biofilm remaining in the wells was quantified as described previously using crystal violet staining [5].

### DNA extraction

DNA was extracted from both initial mixtures (0 h incubation) and end point co-cultures (72 h incubation) using the DNeasy Power Biofilm extraction kit (DNeasy PowerBiofilm, Qiagen, Mississauga, ON) as described previously [5], except that both planktonic (i.e. supernatant of each well) and biofilm fractions (scraped from the bottom of the 96-well plates) were combined for DNA extraction. Extraction controls (reagents only) were included for all batches of DNA extractions and these controls were carried through the PCR, library preparation and sequencing process.

### Amplification of rpoB sequences

Since cpn60 universal primers are degenerate and might introduce amplification bias, we chose rpoB as a target for amplicon sequencing. The rpoB sequences of the nine isolates were aligned using MUSCLE in MegaX, and primers were designed using SnapGene to anneal to perfectly conserved sequences flanking a 353 bp region (corresponding to nucleotides 1441-1793 of the *G. vaginalis* NR001 rpoB gene) that included 29 bp differences between subgroup C and D isolates. Primers were modified with the addition of Illumina adapter sequences (underlined): M_G_RPOBF (5’ - <u>TCG TCG GCA GCG TCA GAT GTG TAA AGA CAG</u> ATG TGC CCA ATC GAA TCC - 3’ and M_G_RPOBR: 5’ - <u>GTC TCG TGG GCT CGG AGA TGT GTA TAA GAG ACA GTC GTG C</u>TC CAA GAA TGG AAT - 3’). Specificity of the primers was confirmed by amplification and sequencing of the target region from isolate NR001. Equal amounts of genomic DNA extracts from all nine isolates were subsequently used as template in rpoB PCR. The amplification products were resolved on an agarose gel and visually inspected to confirm uniform amplification. All products were sequenced to confirm the rpoB sequence of each isolate.

For creation of amplicon sequencing libraries, PCR reactions contained 2 µL template DNA, 1 × PCR buffer (0.2 M Tris-HCl at pH 8.4, 0.5 M KCl), 2.5 mM MgCl_2_, 200 µM dNTP mixture, 400 nM of primers M_G_RPOBF and M_G_RPOBR, 2 U high fidelity platinum Taq DNA polymerase, and water to bring it to a final volume of 50 µL. Control reactions containing no template were included and carried through the library preparation and sequencing steps. PCR was carried out with incubation at 94°C for 30 seconds, 25 cycles of 94°C 30 sec, 60°C for 30 sec, 72°C for 1 min, and final extension at 72°C for 10 min. Purification of the amplified rpoB sequences was performed using NucleoMag magnetic beads (Machery-Nagel, Germany). A bead volume of 40 µl was added per sample to maximize yield of DNA. All samples were run on an agarose gel after the purification step to confirm the size and purity of the PCR product.

### Library generation and amplicon sequencing

Indexing of purified PCR products was performed using the Nextera XT Index kit v2 (Set D) as per manufacturer’s instruction (Illumina Inc, CA, USA). The libraries after indexing were quantified using Qubit dsDNA BR assay kit (Invitrogen, Burlington, Ontario), and indexed amplicons were pooled at an equimolar concentration. PhiX DNA (20% [vol/vol]) was added to the indexed libraries before loading onto the flow cell. Paired-end (2 × 250 bp) sequencing was performed using MiSeq Reagent Nano kit v2 (500 cycles) on an Illumina MiSeq (Illumina Inc., CA, USA). DNA extraction controls and PCR no template controls were carried through library preparation and sequencing.

### Bioinformatics

Because the subgroup C and D rpoB sequences were so distinct, only the R1 data were used in the analysis. Primer sequences were removed using cutadapt [11], and sequences were trimmed for quality using Trimmomatic version 0.32 [12] with a minimum length of 150 bp and quality score of 30. Amplicon sequence variants (ASV) were identified using DADA2 [13] in QIIME 2 [14], which generated a frequency table. ASV sequences were identified by aligning them to a database containing the rpoB sequences of the nine study isolates using watered-BLAST [15].

The proportion of each D isolate in each co-culture was calculated at 0 hours and 72 hours using the equation: R_i_/R_t,_ where R_i_ is the number of sequence reads corresponding to the D isolate in the mixture and R_t_ is the total number of sequence reads obtained for that mixture. The change of proportion of each D isolate in each mixture was calculated using the formula: P_c_ = P_f_/P_i_ −1, where P_c_ is the change of proportions, P_i_ is the proportion of an isolate at 0 h and P_f_ is the final proportion after 72 h. A positive value for P_c_ indicates that the proportional abundance of the D isolate increased over the 72 h incubation period, while a negative value indicates that its proportional abundance in the co-culture decreased. Change of proportion was calculated for each replicate co-culture individually.

### Statistical analysis

To detect differences in the numbers of co-cultures in which C or D had a positive change in proportion, a chi-square test was performed. A chi-square test was performed to determine if the number communities where there was a positive change in proportions for the D isolate was different when its initial proportion was low (P_i_ < 0.3), medium (0.3 < P_i_ < 0.5), or high (P_i_ > 0.5). Change in proportion values for D isolates in co-cultures at different dilutions were compared using Kruskal-Wallis and Dunn’s multiple comparison. Statistical analyses were performed in R version 4.0.1 and GraphPad Prism version 8.

## Results

### Growth of co-cultures

Viability of all isolates prior to mixing and inoculation of 96-well plates was confirmed by observation of growth on blood agar plates. Growth was detected as an increase in OD_595_ in 162/168 co-cultures. Co-cultures containing C2 (NR038) reached a maximum OD at 48 hrs, while the OD of co-cultures containing C1 (NR001) increased between 48 and 72 h (Fig S1). In most cases, as initial culture density decreased (from 10^−1^ to 10^−4^ dilutions), the maximum OD achieved also decreased.

Since the same seven D isolates were included in co-cultures with both C1 and C2, one explanation for the different co-culture growth rates (Fig S1) was that C1 and C2 have different growth rates. To investigate this possibility, C1 and C2 were grown as singletons with different initial densities (Fig S2). C2 grew faster and achieved higher OD values than C1 at all dilutions, reaching its maximum OD (0.2-0.3) by 48 h. The final OD for C2 was not obviously affected by the density of the starter culture. C1, however, was affected by starting density with reduced 72 h OD values with reduced initial density. No growth of C1 was detected in the culture with the lowest initial density (10^−4^ dilution) (Fig S2). Both isolates grew almost exclusively in biofilm (Fig S2).

The seven D isolates were also grown as singletons at varying initial population density to test if the initial population density affects their growth, regardless of the presence of a competitor. Although at the highest population density (10^−1^) D5 and D1 outperformed all other D isolates, D4 reached the highest OD_595_ values in all subsequent dilutions (10^−2^ – 10^−4^) (Fig S3). The other four D isolates: D2, D3, D6 & D7 were consistent in all four dilutions (Fig S3). The optical density of all seven isolates reached an OD_595_ of 0.5 to 1.0 (Fig S3). Except for D5 and D6 at 10^−1^ (Fig S3a), the growth curve of all D isolates at all dilutions plateaued after 48 h. All seven isolates grew exclusively as biofilm (Fig S3e, S3f). Although biofilm formation was appreciably lower at the highest population density (i.e., 10^−1^) compared to more dilute cultures (10^−2^ −10^−4,^ Fig S3f), all subgroup D isolates, in general, formed more biofilm at all dilutions than subgroup C isolates (Fig S2f, Fig S3f).

### rpoB amplicon sequencing

After primer removal and trimming for quality, a total of 1,104,683 reads were available for analysis (average 5225 reads per sample). One of the extraction controls (*n =* 4) yielded 7 reads. All other extraction controls and no template controls (*n* = 4) had 0 reads. Since the no template controls and extraction controls were essentially clean and given the low initial densities of the 10^−4^ dilutions, we decided to include any samples with ≥ 50 reads in the analysis. After removal of samples with no growth based on OD_595_ after 72 h and samples with fewer than 50 reads, 204 samples, including 52 0 h and 152 72 h co-cultures remained for analysis. DADA2 identified twelve ASV sequences, all of which were 98%-100% identical to one of the rpoB sequences of the nine tested isolates.

### Effect of initial abundance of an isolate on the outcome of competition

Read counts corresponding to C and D isolates in each co-culture rpoB amplicon library were used to calculate initial (0 h) and final (72 h) proportions, and the change in proportion of D for each co-culture replicate was calculated. Subgroup D isolates were dominant (>50% of sequence reads) in only 6/52 (11.5%) of the 0 h co-cultures, but at 72 h nearly half (72/152, 47.4 %) were dominated by a D isolate (Fig 2a). Regardless of which isolate was initially more abundant, there was a significantly higher number of co-cultures where proportional abundance of D isolates increased (*n* = 104/152, 68%) compared to those where C isolates had a positive change in proportions (*n =* 48/152, 32%, Chi-square test, p < 0.0001) (Fig 2b).

**Fig 2.**
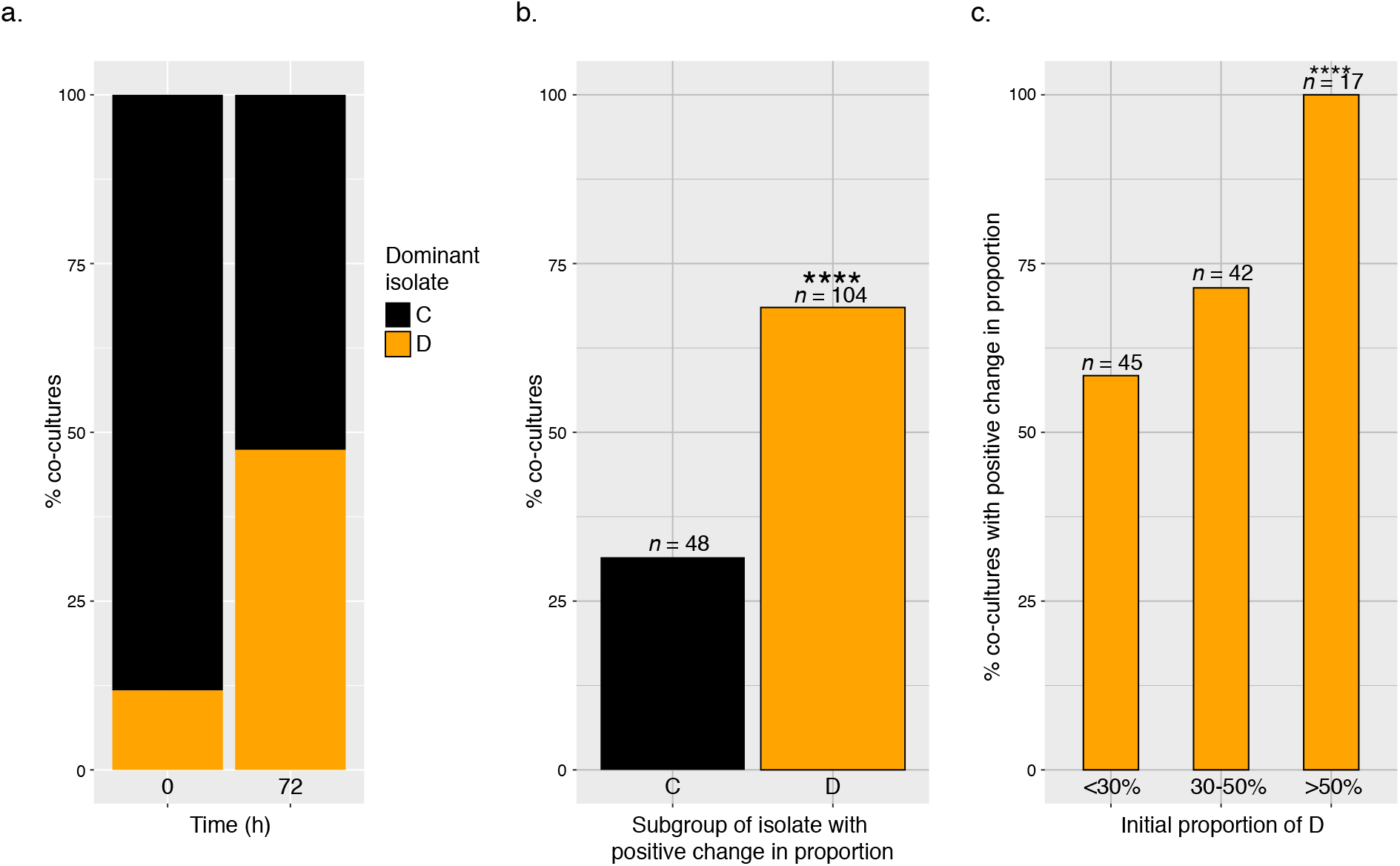
Overall changes in proportion for co-cultures regardless of initial density (a) At 0h most co-cultures were dominated (>50% of sequence reads) by subgroup C, however, subgroup D was dominant in almost half of the 72h co-cultures. (b) Proportion of co-cultures in which C or D had a positive change in proportions. (c) Percent of co-cultures where D increased in proportion when initial proportional abundance was low (<0.3), medium (0.3-0.5) or high (>0.5). Significant effects on change of proportions at varying densities are denoted by asterisks (**** = <0.0001)

Since negative frequency-dependent selection favours rare variants, we tested if initial proportional abundance had an impact on the outcome of D isolates in the competition assay regardless of initial overall population density (dilution). Co-cultures were grouped according to their initial proportions of D (<0.3, n = 45; 0.3 to 0.5, n = 42 or >0.5, n = 17) (Fig 2c). A positive change in proportion of D occurred in all cases where its initial proportional abundance was >0.5, which was significantly higher than when the initial proportional abundance of D was <0.3 or 0.3 – 0.5 (Chi-square test, p < 0.0001), however, at all levels, the proportional abundance of D increased in >50% of cases.

### Effect of starting population density on the outcome of competition

Examination of overall co-culture results demonstrated that in most cases, D isolates experienced an increase in proportion, consistent with negative frequency-dependent selection. If negative frequency-dependent selection is density-dependent, change in proportion values should differ between co-cultures with different initial densities. To determine if the initial population density of co-cultures affects the outcome of competition, seven subgroup D isolates were co-cultured with either of two subgroup C isolates at starting dilutions from 10^−1^ to 10^−4^. All D isolates (D1-D7) had a positive change in proportions at all dilutions when co-cultured with C1 (Fig 3). While D1 and D7 also had positive changes in proportion when co-cultured with C2, the other D isolates had negative changes in proportion at all dilutions (except D6, which was a positive change at the 10^−1^ and 10^−2^ dilutions) (Fig 4).

**Fig 3.**
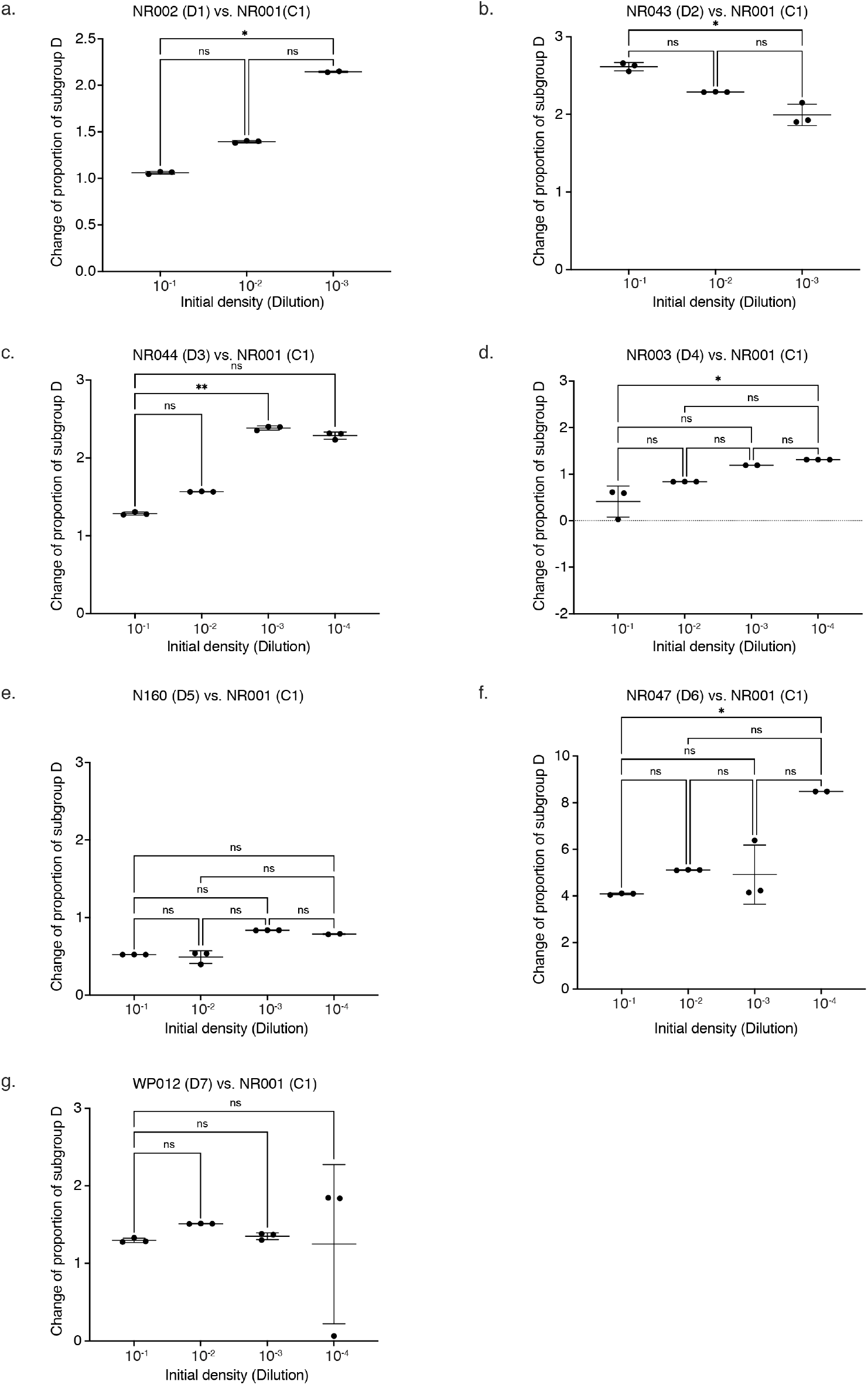
Change in proportion of D1-D7 when co-cultured with C1 at a range of dilutions. Experimental triplicates are plotted with the bars indicating mean with standard deviations. Kruskal-Wallis and Dunn’s multiple comparison was performed to test if the differences of change of proportion at different densities are significant or not. Significant effects on change of proportions at varying densities are denoted by asterisks (* = <0.05, ** = <0.01, *** = <0.001).

**Fig 4.**
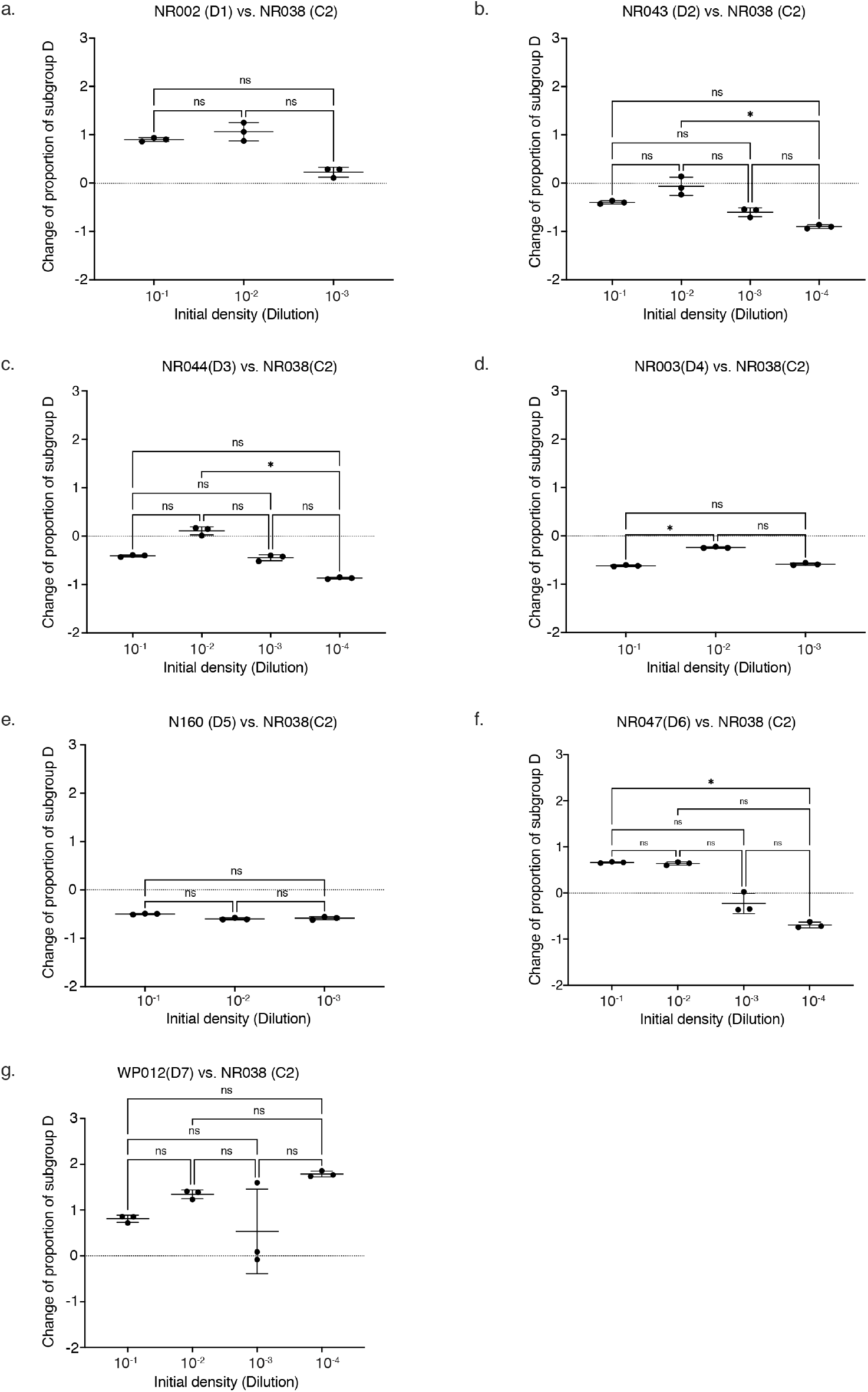
Change in proportion of D1-D7 when co-cultured with C2 at a range of dilutions. Experimental triplicates are plotted with the bars indicating mean with standard deviations. Kruskal-Wallis and Dunn’s multiple comparison was performed to test if the differences of change of proportion at different densities are significant or not. Significant effects on change of proportions at varying densities are denoted by asterisks (* = <0.05, ** = <0.01, *** = <0.001).

The change of proportion of D in eight of the fourteen tested combinations was affected by decreasing density of the starting population either positively (*n* = 4, Fig 3a, 3c, 3d, 3f) or negatively (*n=* 4, Fig 3b, Fig 4b, 4c, 4f) in a statistically significant manner. No effect of initial density was observed for six combinations (Fig 3e, 3g, 4a, 4d, 4e, 4g). All cases where change of proportion of D increased with decreasing density involved co-cultures with C1. In contrast, most (3/4) cases of decreasing change of proportion with decreasing density occurred in co-cultures with C2. These differences in outcomes of co-cultures with C1 and C2 are consistent with the slower growth rate of C1 relative to C2 (Figure S2).

## Discussion

Negative frequency-dependent selection favours rare genotypes [1, 2]. Factors like generalist lifestyle, social cheating, and bet hedging, can contribute to the selection of rare genotypes [8, 9, 16–20]. It has been shown that rarely abundant *Gardnerella* subgroup D are nutritional generalists [8], which may help them to be favoured by negative frequency-dependent selection. However, the fact that subgroup D has not been observed to dominate the vaginal microbiome begs the question: if rare subgroup D is favoured by negative frequency-dependent selection, why do not we see microbiomes dominated by subgroup D more often? The answer to this question probably lies in the mechanisms of negative frequency-dependent selection and that it may be density-dependent [4, 21]. When population density is high, abundant specialists will quickly run out of resources, allowing initially rare generalist populations to expand; however, if the density is lower there are relatively unlimited resources for the rapidly growing specialist, allowing it to maintain numerical dominance (Fig 1). Dynamics of the vaginal microbiome can affect the population density of the vaginal microbiota and the resources available, and hence, may check the expansion of rare species. In this study, we have tested the impact of different initial population densities the outcome of co-cultures of *Gardnerella* strains *in vitro*.

Fourteen combinations of two subgroup C and seven subgroup D isolates were tested. The result of amplicon sequencing showed that in most co-cultures, D isolates had a positive change in proportional abundance regardless of initial proportion (Fig 2), and that initial population density did affect the degree of change in proportion of subgroup D isolates in most cases (Fig 3 &4). In some cases, we observed what would have been predicted for density-dependent negative frequency-dependent selection of subgroup D isolates: a decrease in change in proportion values with decreasing population density (Fig 3b, Fig 4a, 4b, 4c, 4d, 4f).

Increasing change in proportion values with decreasing population density, however, were also observed (Fig 3a, 3c, 3d, 3f). These opposing trends were observed for most D isolates when co-cultured with either C isolate, so it is clear that the effects of density on growth of an isolate in co-culture are determined by both the intrinsic characteristics of the isolate itself but also its competitor as has been reported for wound-colonizing bacteria [22]. Thus, the observed variability of the impact of initial population density on co-cultured *Gardnerella* isolates is not unexpected.

Differential growth rate is an obvious explanation for some of our observations. Our experimental design was based on the premise that subgroup C isolates are faster growing than the D isolates, but the singleton cultures (Fig S2, S3) showed that under the conditions used this was not the case. Both C isolates grew to lower OD values than any of the D isolates, and C1 (NR001) was also negatively affected by the starting density when grown alone (Fig S2). All seven D isolates, however, were fairly consistent when they were grown alone and were not affected appreciably by the lowering of initial population density (Fig S3). It should be noted that we used OD_595_ of the entire well to measure growth and all isolates grew primarily in biofilm (Fig S2f, Fig S3f) and so we cannot rule out the possibility that the higher OD_595_ of the seven D isolates can be partially attributed to greater biofilm biomass.

The relatively poor growth of C1 likely accounts for what was observed in co-cultures since given increasingly more room to grow, the faster grower will have more opportunity to gain in proportional abundance. When the D isolates were co-cultured with C2 with its greater growth rate, the outcomes were markedly different and, in most cases, change of proportion values for D isolates decreased with decreasing initial population density. Exceptions to this pattern were isolates D2 and D7 (Fig 3b, 4b & Fig 3g, 4g) that showed the same results (decreasing or increasing change in proportion values, respectively) when co-cultured with either C isolate, demonstrating that growth rate alone does not determine the effects of population density on co-culture outcome.

Differences of growth while competing for resources can also be affected by spatial organization of the community [23]. Since all the competing species used in this experiment grow almost exclusively as biofilms in the culture conditions used in our study (Fig S2f and Fig S3f), it is probable that the success of each isolate was also influenced by competition for gaining a foothold in biofilms [24].

In this study, the effect of initial population density on competing isolates were conducted in a static culture system, which cannot recapitulate the dynamics of the vaginal microbiome and does not include other members of the vaginal microbiota [6, 25], making it an inadequate model for other mechanisms that may influence the maintenance of rare species. Niche capacity is one such factor that may be important *in vivo* [10, 17, 26]. While one species is the best competitor in its own niche, it may not fare well when invading a neighboring niche. As a result, the numerical abundance of that species is limited to the capacity of its niche. We have shown previously studies that the rare *Gardnerella* species (subgroup D) have minimal overlap with the other *Gardnerella* spp. and are probably occupying a different niche in the vagina [8]. As a result, regardless of its competitive ability, if the capacity of subgroup D’s niche is smaller than one of the more common but less competitive subgroups, its abundance will be limited [9, 10, 26].

It would be naïve to imagine that one ecological mechanism alone can seal the fate of any species in an ecosystem [1, 10]. Taken together, our current study shows that the abundance of species in a mixed *Gardnerella* community can be affected by changing population density, but also highlights the complexity of interacting factors and mechanisms at play. Further advances in our understanding of vaginal microbial community dynamics will be possible with improved model systems for longer-term or continuous culture of consortia in an easily manipulated environment that also allow sufficient replication for robust experimental design.

## Acknowledgements

Thanks to Champika Fernando for excellent technical support and to all members of the Hill Lab for helpful discussions and feedback. No thanks to COVID-19.

## Declarations

### Funding

The research was supported by a Natural Sciences and Engineering Research Council of Canada Discovery Grant to JEH.

### Competing interests

None declared.

### Authors’ contributions

Conceived and designed the study: Salahuddin Khan and Janet E. Hill. Performed the experiments: Salahuddin Khan. Analysed the data: Salahuddin Khan, Janet E. Hill. Wrote and revised the manuscript: Salahuddin Khan, Janet E. Hill.

---

**Fig S1.**
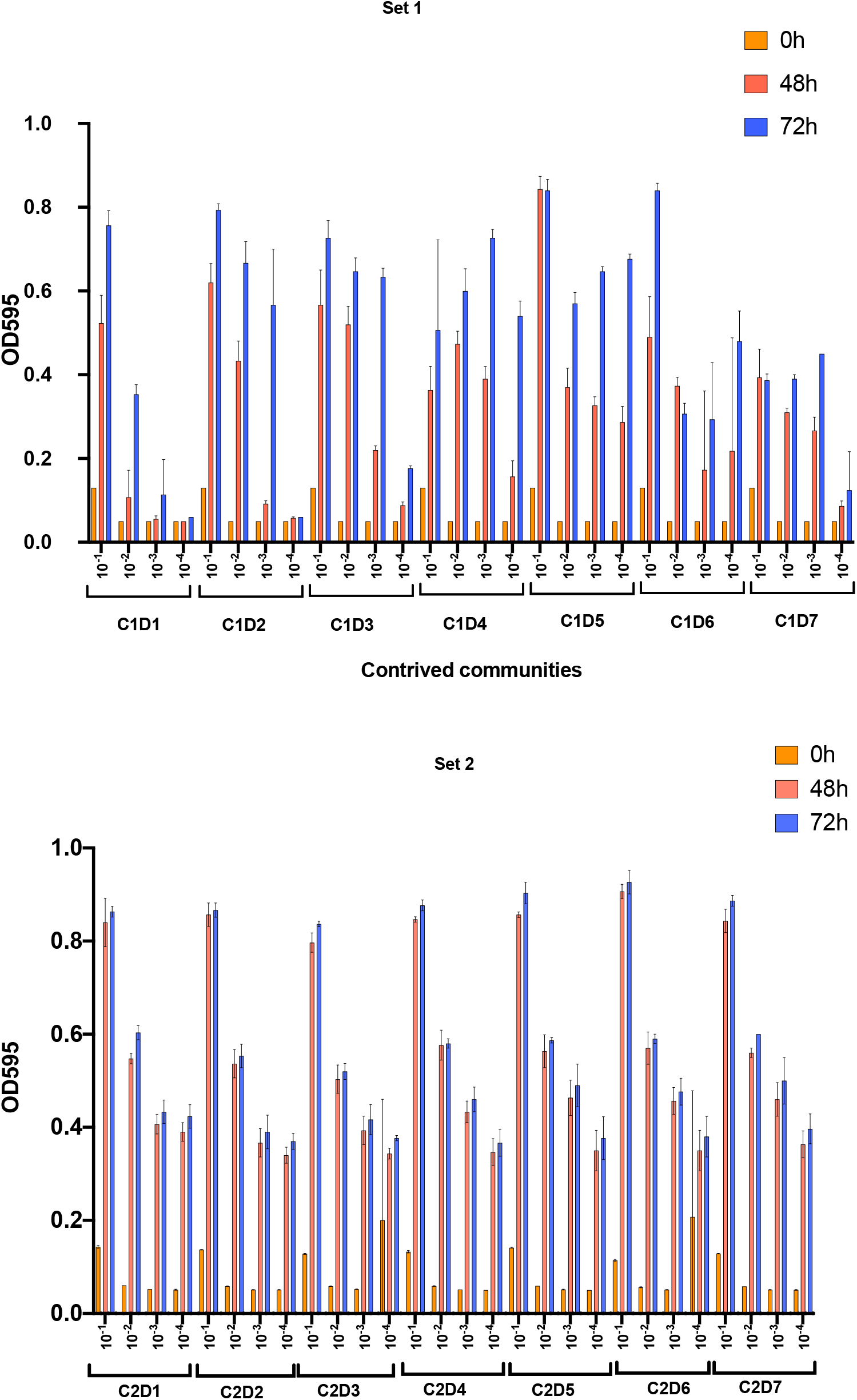
Growth of contrived *Gardnerella* communities at different starting densities based on optical density at 595nm. Values at 48 and 72 h are average of triplicate readings, error bars represent standard deviations of replicates.

**Fig S2.**
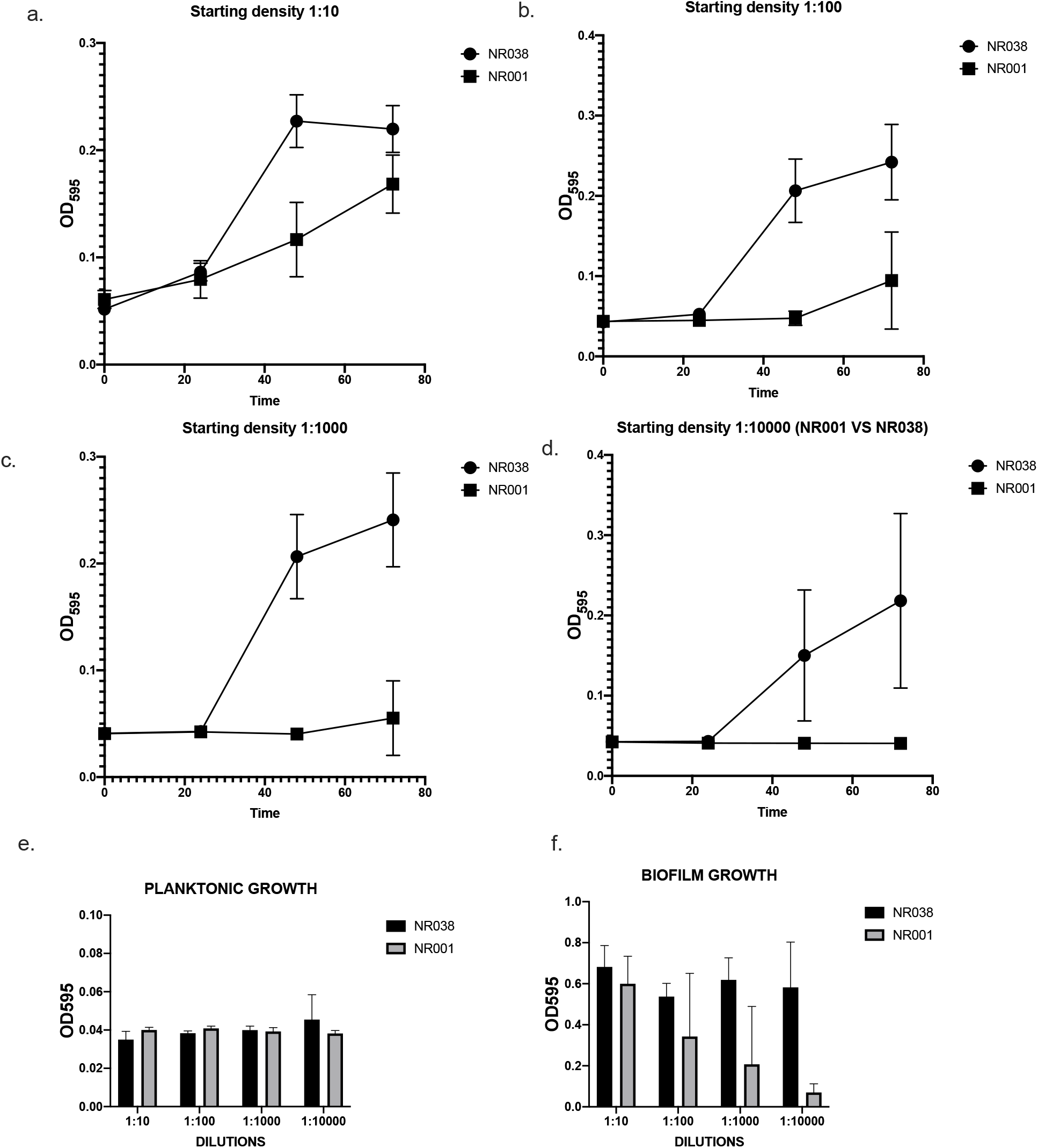
Growth curves of subgroup C isolates NR001 (C1) and NR038 (C2). Subgroup C isolates differ in their growth rate at varying starting density (a-d). Planktonic and biofilm growth of the two isolates after 72h (e, f).

**Fig S3.**
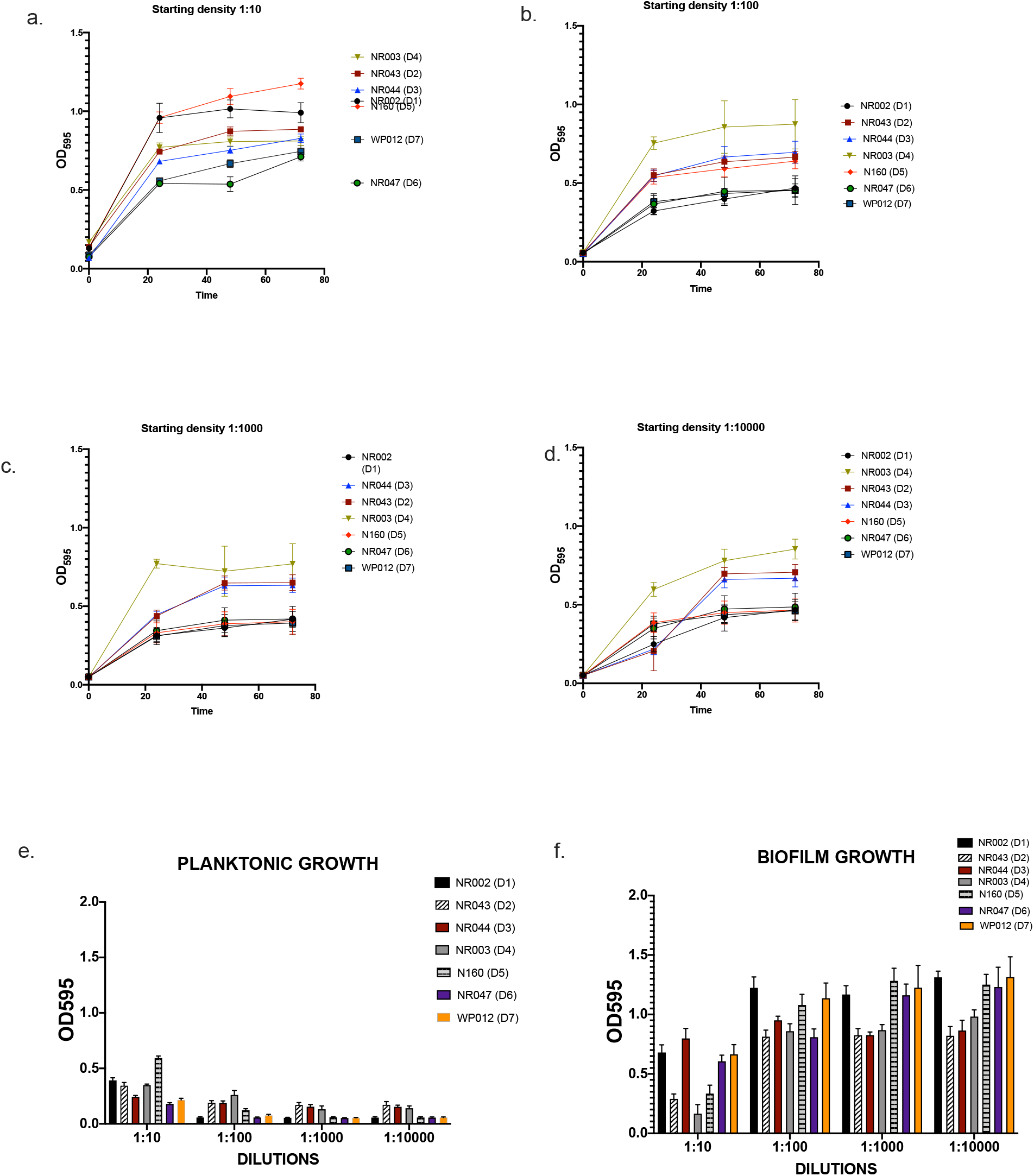
Growth curves of subgroup D isolates. The growth rate of subgroup D is fairly consistent at varying population densities (a-d). All subgroup D isolates grew exclusively as biofilms (e-f).

## Notes

### Competing Interest Statement

The authors have declared no competing interest.

